# A novel gonorrhea vaccine composed of MetQ lipoprotein formulated with CpG shortens experimental murine infection

**DOI:** 10.1101/2020.06.19.161646

**Authors:** Aleksandra E. Sikora, Carolina Gomez, Adriana Le Van, Benjamin I. Baarda, Stephen Darnell, Fabian G. Martinez, Ryszard A. Zielke, Josephine A. Bonventre, Ann E. Jerse

## Abstract

Bacterial surface lipoproteins are emerging as attractive vaccine candidates due to their biological importance and the feasibility of their large-scale production for vaccine manufacturing. The global prevalence of gonorrhea, resistance to antibiotics, and serious consequences to reproductive and neonatal health necessitate development of effective vaccines. Reverse vaccinology identified the surface-displayed L-methionine binding lipoprotein MetQ (NGO2139) and its homolog GNA1946 (NMB1946) as gonococcal and meningococcal vaccine candidates, respectively. Here, we assessed the suitability of MetQ for inclusion in a gonorrhea vaccine by examining MetQ conservation, its function in *Neisseria gonorrhoeae* (*Ng*) pathogenesis, and its ability to induce protective immune responses using a female murine model of lower genital tract infection. In-depth bioinformatics, phylogenetics and mapping the most prevalent *Ng* polymorphic amino acids to the GNA1946 crystal structure revealed remarkable MetQ conservation: ~97% *Ng* isolates worldwide possess a single MetQ variant. Mice immunized with rMetQ-CpG (*n*=40), a vaccine containing a tag-free version of MetQ formulated with CpG, exhibited robust, antigen-specific antibody responses in serum and at the vaginal mucosae including secretory IgA. Consistent with the activity of CpG as a Th1-stimulating adjuvant, the serum IgG1/IgG2a ratio of 0.38 indicated a Th1 bias. Combined data from two independent challenge experiments demonstrated that rMetQ-CpG immunized mice cleared infection faster than control animals (vehicle, *p*<0.0001; CpG, *p*=0.002) and had lower *Ng* burden (vehicle, *p*=0.03; CpG, *p*<0.0001). We conclude rMetQ-CpG induces a protective immune response that accelerates bacterial clearance from the murine lower genital tract and represents an attractive component of a gonorrhea subunit vaccine.

## INTRODUCTION

Vaccines against infectious diseases have indisputably transformed human health. There is a huge need for continuous efforts to develop vaccines against challenging bacterial infections caused by drug-resistant and/or immunoregulating pathogens. Recently, surface-exposed lipoproteins have garnered interest for their utility in subunit vaccines against disease-causing microbial pathogens, including *Clostridioides difficile*, Dengue virus, *Neisseria meningitidis* (*Nm*) serogroup B, *Burkholderia cepacia*, and Human Papilloma Virus [1–3]. Lipoproteins play myriad roles in bacterial physiology, which range from acquiring nutrients, contributing to virulence, establishing and maintaining cell envelope homeostasis, and interacting with host cells [4]. During infection, lipoproteins may also act as Toll-like receptor (TLR)2 agonists and contribute to an immune response against the pathogen [1, 5–7]. Characterized by an invariant cysteine residue that is post-translationally modified with two to three acyl chains, lipoproteins are tethered to either the cytoplasmic or outer membrane in Gram-negative bacteria through this lipid anchor and may face either the periplasmic space or the extracellular environment [7–10]. Biochemically, lipoproteins are frequently hydrophilic, which facilitates their production in a heterologous host and yields large scale quantities of soluble and natively folded proteins in a cost-effective manner, which are important considerations for vaccine commercialization. Subunit antigens, such as lipoproteins, are also appealing candidates for vaccine development due to their safety and reduced chance of side effects [11, 12].

In concert with gonorrhea vaccine development endeavors, here we focused on the pre-clinical evaluation of MetQ, which is a surface-displayed lipoprotein that elicits strongly bactericidal antibodies [13, 14], as a component of a gonorrhea subunit vaccine. The causative organism of this sexually transmitted infection, *Neisseria gonorrhoeae* (*Ng*), is a Gram-negative diplococcus that exclusively plagues humankind, yielding about 87 million new cases annually across the globe [15]. In the United States, it remains the second most commonly reported notifiable disease and the number of cases has risen steadily since the historic low in 2009, increasing by 82.6% (a total of 583,405 reported cases) in 2018 [16]. The consequences of gonorrhea and other sexually transmitted infections have devastating health and psychological impacts that profoundly affect quality of life [17]. Clinical presentations of gonorrhea vary between infection site and gender and include cervicitis, urethritis, proctitis, conjunctivitis, or pharyngitis. Women tend to have more frequent asymptomatic urogenital infections and experience serious long-term reproductive health problems including endometritis, pelvic inflammatory disease, pregnancy complications and infertility [18]. Disseminated gonococcal infection occurs in 0.5-3% patients and can lead to musculoskeletal manifestations, such as suppurative arthritis, or a distinct syndrome that includes tenosynovitis, skin lesions, and polyarthralgia [19–21]. Neonates infected during vaginal delivery most typically develop ophthalmia neonatorum, which can lead to blindness if left untreated. Localized infections of other mucosal surfaces can also occur [22]. The grave nature of gonorrhea is further exacerbated by augmenting human immunodeficiency virus (HIV) infectivity and susceptibility to HIV [23]. In the United States, patients presenting with uncomplicated *Ng* infections are given a single dose of injectable ceftriaxone and oral azithromycin in compliance with treatment guidelines from the Centers for Disease Control and Prevention [24]. Alarmingly, however, these antimicrobials are rapidly losing their effectiveness [25–27]. Continued efforts to develop a gonorrhea vaccine are critical.

The MetQ immunogen investigated herein was identified by proteomics and genomics-based reverse vaccinology antigen discovery programs applied to *Ng* [13, 28, 29] and *Nm* [13, 28–30], respectively. Using high-throughput proteomics and immunoblotting analyses, we previously reported that MetQ is ubiquitously expressed in diverse *Ng* isolates [13, 28, 29, 31], in host-relevant growth conditions [13] and in naturally released outer membrane vesicles [13, 28]. In addition to a traditional lipoprotein signal peptide, MetQ contains a NlpA domain homologous to the respective domain in the *E. coli* methionine binding protein MetQ, NlpA [14, 32]. Indeed, *Ng* MetQ binds L-methionine with nanomolar affinity. Besides its function in methionine import, MetQ impacts *Ng* adhesion and invasion of epithelial cells and bacterial survival in primary monocytes, macrophages, and human serum [14].

In this report, we employed bioinformatics to assess MetQ conservation on a large scale, examined its importance in *Ng* pathogenesis using a mouse infection model, and evaluated its *in vivo* efficacy against *Ng* challenge when formulated with a T helper (Th) 1 response-inducing adjuvant (CpG). Our study demonstrates that rMetQ-CpG induces a robust systemic and vaginal humoral immune response and significantly shortens experimental gonococcal infection. Further, we reveal that MetQ is exceptionally conserved and propose its inclusion in a broad-spectrum gonorrhea vaccine.

## MATERIALS AND METHODS

### Study Design

#### Sample size and number of replicates

Twenty mice per experimental group were immunized in each of two biologically independent immunization/challenge studies. The number of animals was chosen based on our extensive experiences with mouse studies and was selected using power predictions based on the observation that only ~75% of mice will be in the correct stage of the estrous cycle at the time of challenge. Mice were randomly assigned to treatment groups and data points from all individual animals were included in analyses. No outliers were defined or excluded. For all experiments, statistical significance and number of replicates are indicated in the figure legends, the main text, or the following sections.

### Bacteria and Culture Conditions

*Ng* strain FA1090 (serum-resistant, PorB1B) was originally isolated from a female with disseminated gonococcal infection [33] and was used in the studies. The isogenic knockout *ΔmetQ* and complementation strain Δ*metQ/P_lac_::metQ* were constructed previously [13].

*Ng* strains were routinely cultured on GC agar (GCB; Difco) containing Kellogg’s Supplement I and 12.5 μM ferric nitrate in a 5% CO_2_ atmosphere at 37°C for 18-20 h, as described [34]. After passage on GCB, transparent and non-piliated colonies were cultured in gonococcal base liquid (GCBL) medium supplemented with Kellogg’s supplement I in 1:100 and 12.5 μM ferric nitrate [35, 36]. Viability of *Ng* strains was evaluated under conditions mimicking different host microecological niches as described [37].

To isolate *Ng* from facultatively anaerobic commensal flora from mouse vaginal swabs, GCB with vancomycin, colistin methane sulfonate, nystatin, trimethoprim (VCNT) and 100 μg/mL streptomycin (VCNTS) and heart infusion agar (HIA) were used, respectively.

*E. coli* strains MC1061 and BL21(DE3) were used as a host during molecular cloning procedures and for overexpression of rMetQ, respectively. *E. coli* were maintained on Luria-Bertani (LB) agar or cultured in LB broth.

Antibiotics were used in the following concentrations: for *Ng:* kanamycin 40 μg/mL, streptomycin 100 μg/mL; for *E. coli:* kanamycin 50 μg/mL, carbenicillin 50 μg/mL.

### Cloning and Purification of rMetQ

To generate a vector encoding recombinant variant of MetQ (rMetQ) with a N-terminal 6×Histidine-tag followed by the Tobacco Etch Virus (TEV) protease recognition site, the *ngo2139* chromosomal region excluding coding sequence for the initial 34 amino acids and the stop codon was amplified with primers rMetQ-F 5’GATATCCCATGGAAAAAGACAGCGCGCCCG3’ and rMetQ-R 5’GGATCCAAGCTTTTTGGCTGCGCCTTCATTCC3’. Obtained PCR product (531 bp) and pRSF-NT vector were digested with NcoI-HF and HindIII-HF (New England BioLabs) and ligated using T4 DNA ligase (New England BioLabs). The obtained pRSF-NT-rMetQ DNA was sequenced at the Center for Genome Research and Biocomputing at Oregon State University.

To purify rMetQ, overnight cultures of *E. coli* BL21(DE3) carrying pRSF-NT-rMetQ were back-diluted (1:100) in 3 L of LB broth and incubated with aeration at 37°C until OD_600_ of ~0.5. Protein overproduction was induced by addition of isopropyl β-D-1-thiogalactopyranoside (IPTG) at a final concentration of 0.1 mM. Cultures were incubated for additional 3 h, harvested by centrifugation, and pelleted cells were suspended in lysis buffer (20 mM Tris-HCl pH 8.0, 500 mM NaCl, 10 mM imidazole) supplemented with a Complete, EDTA-free Protease Inhibitor Cocktail tablet (Roche). Cell lysis was accomplished by passaging six times through a French pressure cell at 12,000 psi. Cell debris and unbroken cells were removed by centrifugation and supernatants were passed through a 0.22 μm filter unit (VWR International). Soluble protein suspension was loaded onto the Bio-Scale Mini Profinity IMAC Cartridge (5 mL; Bio-Rad Laboratories) connected to the NGC Scout Chromatography system (Bio-Rad Laboratories). Unspecific and weakly bound proteins were removed from the column with 8 volumes of lysis buffer mixed with 12% of elution buffer (20 mM Tris-HCl pH 8.0, 500 mM NaCl, 250 mM imidazole). The proteins were eluted with 5 volumes of elution buffer. Fractions containing rMetQ were combined and concentrated with a Vivaspin 20 MWCO 5000 (GE Healthcare), according to the manufacturer’s recommendations. Concentrated samples were loaded onto Hi Load 16/600 Superdex 75 pg column (GE Healthcare) connected to the NGC Scout Chromatography system and running buffer composed of 20 mM Tris pH 8.0, 500 mM NaCl. Fractions containing rMetQ were combined and concentrated using a Vivaspin 20 device. The C-terminal 6×Histidine-tag was cleaved by mixing with TEV protease (20:1 w:w ratio) in 500 μL of cleavage buffer (0.5 M Tris-HCl pH 8.0, 5 mM EDTA, 1 mM dithiothreitol, and 500 μL of Ni-NTA agarose (Qiagen) equilibrated with cleavage buffer. Following overnight incubation at room temperature, the cleavage mixture was loaded onto a 5 mL polypropylene column (ThermoScientific) and supernatant containing a tag-free rMetQ protein was collected.

To remove residual co-eluted TEV protease, sample was mixed with 500 μL of Ni-NTA agarose equilibrated with cleavage buffer, incubated for 1 h with rotation at 4°C, and loaded onto a 5 mL polypropylene column. Collected supernatant containing rMetQ was applied onto a PD-10 column (GE Healthcare) for buffer exchange (PBS, 5% glycerol), and purified rMetQ was aliquoted and stored at −80°C.

### Competitive infections

Competitive infections were performed essentially as described [34] in female BALB/c mice treated with Premarin (0.5 mg, days −2, 0 and +2) and the same antibiotic regimen used for the challenge experiments. Two days after estrogen treatment was initiated (d0), groups of mice (*n* = 5-7 mice/group) were inoculated with saline suspensions containing WT mixed with similar numbers of the Δ*metQ* mutant or the complementation strain Δ*metQ/P_lac_::metQ* (total dose: 1×10^6^ CFUs/mouse). Vaginal swabs were collected on days 1, 3 and 5 post-inoculation, suspended in 1 mL of GCBL with 0.05% saponin and serially diluted 10^−1^ to 10^−4^. Similar volumes of undiluted and diluted suspensions were inoculated onto GCB supplemented with streptomycin (total CFUs) or GCB containing streptomycin and kanamycin (CFUs of Δ*metQ* mutant or Δ*metQ*/*P_lac_*::*metQ*). The number of WT bacteria was determined by subtracting the number of CFUs recovered on plates with streptomycin and kanamycin from the number recovered on solid media with streptomycin. Relative fitness was determined by calculating the CI using the equation CI = [mutant CFU (output)/WT CFU (output)]/[mutant CFU (input)/WT CFU (input)]. The limit of detection of 1 CFU was assigned for a strain that was not recovered from an infected mouse. A CI of <1 or > 1 indicates that the mutant is less or more fit than the WT strain, respectively. Two independent experiments were conducted and the results were similar. Statistical analysis was performed using Kruskal-Wallis Dunn’s multiple comparison tests to compare CIs between the Δ*metQ/WT* and Δ*metQ/P_lac_::metQ/*WT.

### Immunization and challenge studies

Female BALB/c mice (3-4 weeks old, Charles River Laboratories, NCI BALB/c strain) were immunized by subcutaneous injection of 20 μg of rMetQ mixed with 80 μL of Titermax Gold (Invivogen) (total volume, 150 μL) [day 0 (d0)] followed by three intranasal (IN) boosts on days 14, 24 and 35 consisting of 20 μg of antigen and 20 μg of CpG 1826 (InvivoGen; total volume 40 μL) given in 2, 10 μL volumes per nare, 5 min apart. Control groups received Titermax Gold (d0) and CpG only (d14, 24, 35) or PBS by the same routes. Venous blood and vaginal washes were collected on days 34 and venous blood only on day 49. For vaginal washes, 30 μL of PBS were pipetted in and out of the vagina 3-5 times and pooled. Blood and vaginal washes were centrifuged for 5 min in a microfuge, and the serum fraction and vaginal wash supernatants were stored at −20°C until further analysis. Three weeks after the final immunization, mice in the diestrus or anestrus stage of the reproductive cycle were identified by cytological examination of vaginal smears and implanted with a slow-release 6.5 mg 17β-estradiol pellet (Innovative Research of America) as described [38]. Mice were treated with streptomycin, vancomycin and trimethoprim to suppress the overgrowth of commensal flora that occurs under the influence of estradiol [39]. Two days after pellet implantation, mice were challenged with 10^6^ colony-forming units (CFU) of *Ng* FA1090. Vaginal swabs were collected every other day for 7 days and suspended in 1 mL of GCBL. Swab suspensions were quantitatively cultured for *Ng* using the SpiralTek plater (AP5000) and Qcount™ model 530. A portion of the vaginal swab was inoculated onto HIA to monitor the presence of potentially inhibitory commensal flora and smeared on glass slides for assessment of polymorphonuclear leukocytes (PMNs) influx as described [38]. The percentage of mice with positive cultures at each time point was plotted for each experimental group as a Kaplan Meier curve and analyzed by the Log Rank test. To assess the bacterial burden over time, the mean AUC (log_10_ CFU versus time) was calculated for each mouse. The geometric means under the curve were compared between the groups using the Kruskal-Wallis nonparametric rank sum test and pairwise comparisons between groups were performed with Dunn’s post hoc test as described [40].

### Ethics Statement

Animal experiments were conducted at the Uniformed Services University of the Health Sciences (USUHS) according to the guidelines of the Association for the Assessment and Accreditation of Laboratory Animal Care under a protocol # MIC16-488 that was approved by the University’s Institutional Animal Care and Use Committee. The USUHS animal facilities meet the housing service and surgical standards set forth in the “Guide for the Care and Use of Laboratory Animals” NIH Publication No. 85-23, and the USU Instruction No. 3203, “Use and Care of Laboratory Animals”. Animals are maintained under the supervision of a full-time veterinarian. For all experiments, mice were humanely euthanized by the laboratory technician upon reaching the study endpoint using a compressed CO_2_ gas cylinder in LAM as per the Uniformed Services University (USU) euthanasia guidelines (IACUC policy 13), which follow those established by the 2013 American Veterinary Medical Association Panel on Euthanasia (http://www.usuhs.mil/usuhs_only/lam/lamws/euth.html).

### ELISAs

Titers of antigen-specific total IgG, IgG1, IgG2a and IgA were measured in serum and vaginal washes by ELISA. Standard antigen titration with serially diluted pooled serum from each test group was used to identify the lowest concentration of rMetQ that gave a linear curve above background when plotted against antiserum dilution. Round bottom high-binding 96 well microtiter plates (Nunc) were coated with 0.5 μg of rMetQ suspended in phosphate buffered saline (PBS) and were blocked with PBS supplemented with 3% Bovine Serum Albumin. Serum or vaginal wash samples were serially diluted in PBS supplemented with 0.05% Tween (PBST) and added to each well. The wells were washed with PBST and incubated with secondary antisera: anti-mouse total IgG, anti-mouse IgG1, anti-mouse IgG2a, and anti-mouse IgA (Southern Biotech) conjugated to horse radish peroxidase (HRP). Wells were washed and reactions were developed using TMB Peroxidase EIA Substrate (BioRad). End-point titers were determined using the average reading of 8 wells incubated with secondary but no primary antibody plus 3 and 2 standard deviations as baseline for serum and vaginal secretions; respectively. For statistical analysis, Kruskal-Wallis with Dunn’s multiple comparisons test was applied on non-transformed arithmetic data, with *p* values of <0.5 considered significant.

### MetQ bioinformatics analyses

Polymorphism, phylogenetic, and allele mapping analyses were performed as described by [32], with the following exceptions. Briefly, the nucleotide sequence of *metQ* (*ngo2139*) was used to query the public *Neisseria* multilocus sequence typing database (*Neisseria* pubMLST; [41]; https://pubmlst.org/neisseria/) to identify *metQ* alleles and nucleotide polymorphic sites across 4,421 isolates of *N. gonorrhoeae* for which *metQ* sequence data exist, as well as among all *Neisseria* isolates present in the database (21,798 isolates with *metQ* sequence data as of March 26, 2020). Translated amino acid sequences were aligned with Muscle in Geneious Prime 2020.1.2. Alignments were used to generate a Neighbor-Joining phylogenetic tree using the Jukes-Cantor genetic distance model. Amino acid polymorphisms were mapped to crystal structures of the *Nm* MetQ homolog, GNA1946. The amino acid sequences of structures solved with L-methionine [6OJA; [42]] or in the substrate-free conformation [6CVA; [42]] were aligned against all *Ng* MetQ amino acid sequences as above. Polymorphic sites were identified and their prevalence was calculated by dividing the number of isolates associated with each polymorphism by the total number of isolates. The single polymorphic site found in >1% of the global *Ng* population was then mapped to the crystal structure using PyMOL (https://pymol.org/2/).

### Etest antimicrobial sensitivity assessments

The susceptibility of WT FA1090, isogenic knockout Δ*metQ*, and complementation strain Δ*metQ/P_lac_::metQ* to seven antimicrobial compounds was assessed by Etests as previously [37], according to manufacturer’s instructions. Briefly, non-piliated colonies of each strain were suspended in GCBL to a turbidity equivalent to that of a 0.5 McFarland standard and spread on 150 mm tissue culture dishes filled with 50 mL GCB to a thickness of 4 mm. Etest strips were placed on the agar surface and plates were cultured for ~22 h, at which point the MICs were determined. Sensitivity assessments were performed three times, and MIC values that repeated at least twice are reported.

### Isolation of *Ng* crude cell envelope fractions

Total cell envelope (CE) fractions were prepared from mid-logarithmic cultures of WT *Ng* FA10190 by sonication and ultracentrifugation. Briefly, bacteria were cultured in supplemented GC broth to an A_600_ of 0.6, collected, resuspended in PBS and disrupted by sonication. Cell debris and remaining intact cells were removed by centrifugation. Total cell envelope fraction was obtained from crude cell lysates by ultracentrifugation, resuspended in PBS and frozen at −20°C. Protein concentration was measured using a NanoDrop Spectrophotometer (ND-1000).

### Immunoblotting and SyproRuby staining

Samples of purified MetQ (1 μg), crude cell envelope fractions (CE; 1 μg), or vaginal washes (normalized based on CFUs) were fractionated by SDS-PAGE, transferred onto nitrocellulose membranes and detected by immunoblotting as described previously [13]. Protein concentration was measured using the DC Protein Assay (BioRad). The immunoblotting analysis of MetQ expression during murine infection was performed using polyclonal rabbit antisera against MetQ [13], and secondary anti-goat anti-rabbit HRP conjugated antibodies (Bio-Rad). For experiments assessing specificity of murine sera and vaginal washes, membranes were blocked overnight in PBST supplemented with 5% non-fat dry milk and incubated with antiserum (1:5,000) or vaginal washes (1:500) from test or control mice followed by goat anti-mouse IgG (BioRad) or IgA (SouthernBiotech) conjugated to HRP. Washes between incubations were performed with PBST. Membranes were developed using ECL Prime (Amersham) and ImageQuant™ LAS 4000 (GE Healthcare). Proteins were visualized with SyproRuby (BioRad) per manufacturer’s recommendations.

### Densitometry analysis

The expression of MetQ during WT FA1090 infection in female BALB/c mice and intensities of protein bands detected in serum and vaginal specificity studies were determined by densitometry using FIJI software [43]. The amount of MetQ on day 1 post-infection was arbitrarily set to 1 and the protein abundance on days 3 and 5 is expressed relative to the MetQ cellular pool detected on day 1.

### Statistical analysis

GraphPad Prism was used for all statistical analyses. The built-in *t*-test was utilized to determine statistically significant differences between experimental results with the exception of animal studies and ELISA, which were analyzed as described above. A confidence level of 95% was used for all analyses.

## RESULTS

### MetQ is highly conserved among *Neisseria*

We previously analyzed MetQ polymorphic sites between 36 genetically and temporally diverse *Ng* isolates including the 2016 World Health Organization reference strains. MetQ had only 2 (0.23%) and 0 variations at the DNA and protein levels, respectively, and was the most conserved vaccine antigen compared to LptD, BamA, TamA, and NGO2054 [13]. To further explore the feasibility of MetQ as an effective vaccine target, we first comprehensively examined the prevalence of this antigen and its sequence variability using all available *Neisseria* genome sequences deposited to the PubMLST database and their predicted amino acid sequences. Overall, these investigations corroborated that MetQ is highly conserved, with 22 alleles among 4,421 *Ng* isolates, which accounted for 49 nucleotide polymorphic sites and 17 amino acid polymorphisms (Fig. 1a). Remarkably, a single amino acid sequence, represented by 10 of the 22 alleles, accounts for nearly 97% of *metQ* sequence variation across the *Ng* isolates deposited in the database (Table S1). A phylogenetic analysis indicated that all of the *metQ* alleles were closely related, with the exception of allele 41, which formed an outgroup (Fig. 1b). Polyclonal rabbit antisera elicited by recombinant MetQ (rMetQ) representing allele 8 (*Ng* FA1090) previously detected MetQ in whole-cell lysates of diverse *Ng* isolates. Together, these data confirm the ubiquitous nature of this antigen and the conservation of immunogenic epitopes [13].

**Figure 1.**
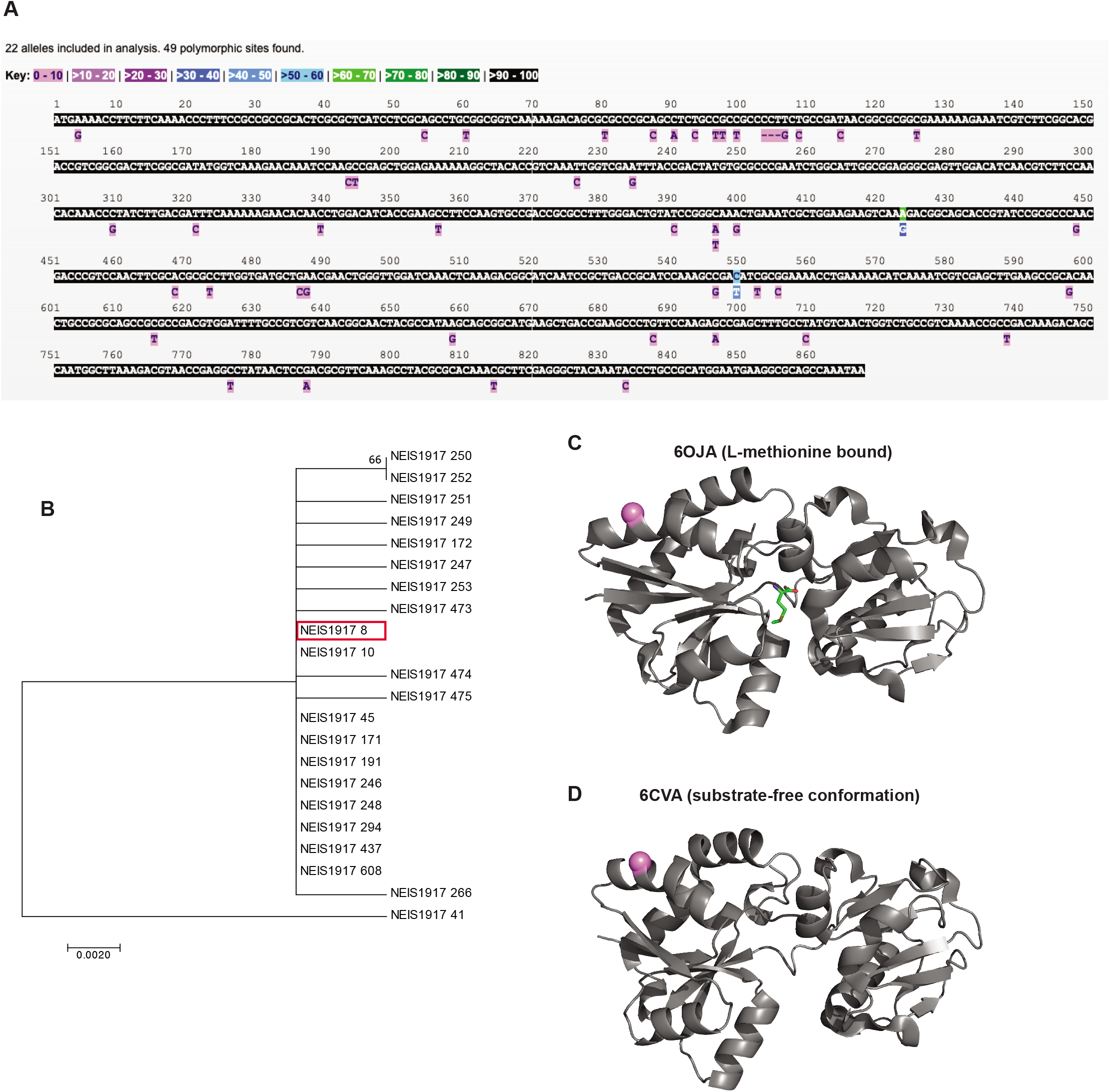
Conservation of *Ng* MetQ and mapped amino acid polymorphic sites. (**A**) Nucleotide polymorphic sites in the *metQ* locus (NEIS1917) across 4,421 *N. gonorrhoeae* isolates with *metQ* sequence data deposited into the *Neisseria* PubMLST database. (**B**) A Neighbor-Joining phylogenetic tree was constructed for *metQ* alleles in Geneious using the Jukes-Cantor genetic distance model. The FA1090 allele (8) is highlighted in red. (**C**, **D**) The single *Ng* MetQ amino acid polymorphism found in >1% of the global *Ng* population was mapped to crystal structures of *Nm* MetQ using PyMol. The light purple sphere indicates position 65, which is polymorphic in 2.65% of isolates. Structures collected with L-methionine (PDB ID: 6OJA; **C**) or or in the substrate-free conformation (PDB ID: 6CVA; **D**) are presented. L-methionine is shown in green in the MetQ active site in the L-methionine-bound structure.

A subsequent broader investigation (*n*=21,798) of other *Neisseria* isolates with *metQ* sequence information revealed 415 *metQ* alleles, which accounted for 645 and 194 nucleotide and amino acid polymorphic sites, respectively (Fig. S1). Similar to *Ng*, *metQ* alleles across *Neisseria* were closely related, with the exception of 13 sequences which clustered separately from the rest of the tree (Fig. S2).

To facilitate future structural vaccinology efforts, we next mapped the most prevalent gonococcal polymorphic amino acids to *Nm* MetQ crystal structures both with and without L-methionine in the binding site (6OJA and 6CVA, respectively [42]). Of the 11 polymorphisms encompassed by the crystal structures, only one site (A65) diverged from the most common MetQ variant in 2.65% of isolates worldwide. The remaining polymorphisms were found in less than 1% of *Ng* MetQ amino acid sequences. (Fig. 1c, D; Supplemental Videos 1-4). Six polymorphic sites outside of the crystal structure [the crystal structure sequence starts at site 44, and the polymorphisms are at sites 2, 27, 33, 35 (gap site), 36, and 42 are also found in fewer than 1% of *Ng* isolates.

These investigations, performed for the first time on a large scale, demonstrate the exceptionally high level of MetQ conservation and strongly support its inclusion in a broad-spectrum gonorrhea vaccine.

### MetQ is expressed *in vivo* but does not confer a detectable advantage during competitive murine infection

Prokaryotic lipoproteins play versatile functions ranging from cell envelope stability to nutrient acquisition, substrate binding for ABC transporter systems, modulation of the host immune system, signal transduction, and virulence [4, 8]. In addition to its predicted role in bacterial physiology as a methionine transporter, MetQ may contribute to gonococcal pathogenesis based on the report that a *Ng DmetQ* mutant was attenuated during exposure to primary monocytes and activated macrophages and was less able to adhere to and invade human cervical epithelial cells [14]. These observations prompted us to further explore phenotypes associated with deletion of *metQ* using previously constructed a null mutant in *ngo2139* (Δ*metQ*) and its complemented mutant Δ*metQ/P_lac_::metQ* in *Ng* FA1090 [13]. We first examined whether complete elimination of MetQ affects cell envelope homeostasis by exposing bacteria to seven antibiotics with different mechanisms of action using Etest assays. These studies showed that *Ng* lacking MetQ had the same susceptibility as the parental strain (Table S2), suggesting that this lipoprotein has no significant function in cell envelope stability. Subsequently, we investigated the role of MetQ during conditions that mimic environmental microniches in the host by exposing WT, Δ*metQ*, and the complementation strain Δ*metQ/P_lac_::metQ* to iron limitation, normal human serum, anaerobiosis, and a combination of iron limitation and anoxia. Loss of MetQ did not significantly alter bacterial viability as assessed by comparing the number of colony forming units (CFU) of the mutant and WT strains following exposure to the conditions examined (Fig. 2a).

**Figure 2.**
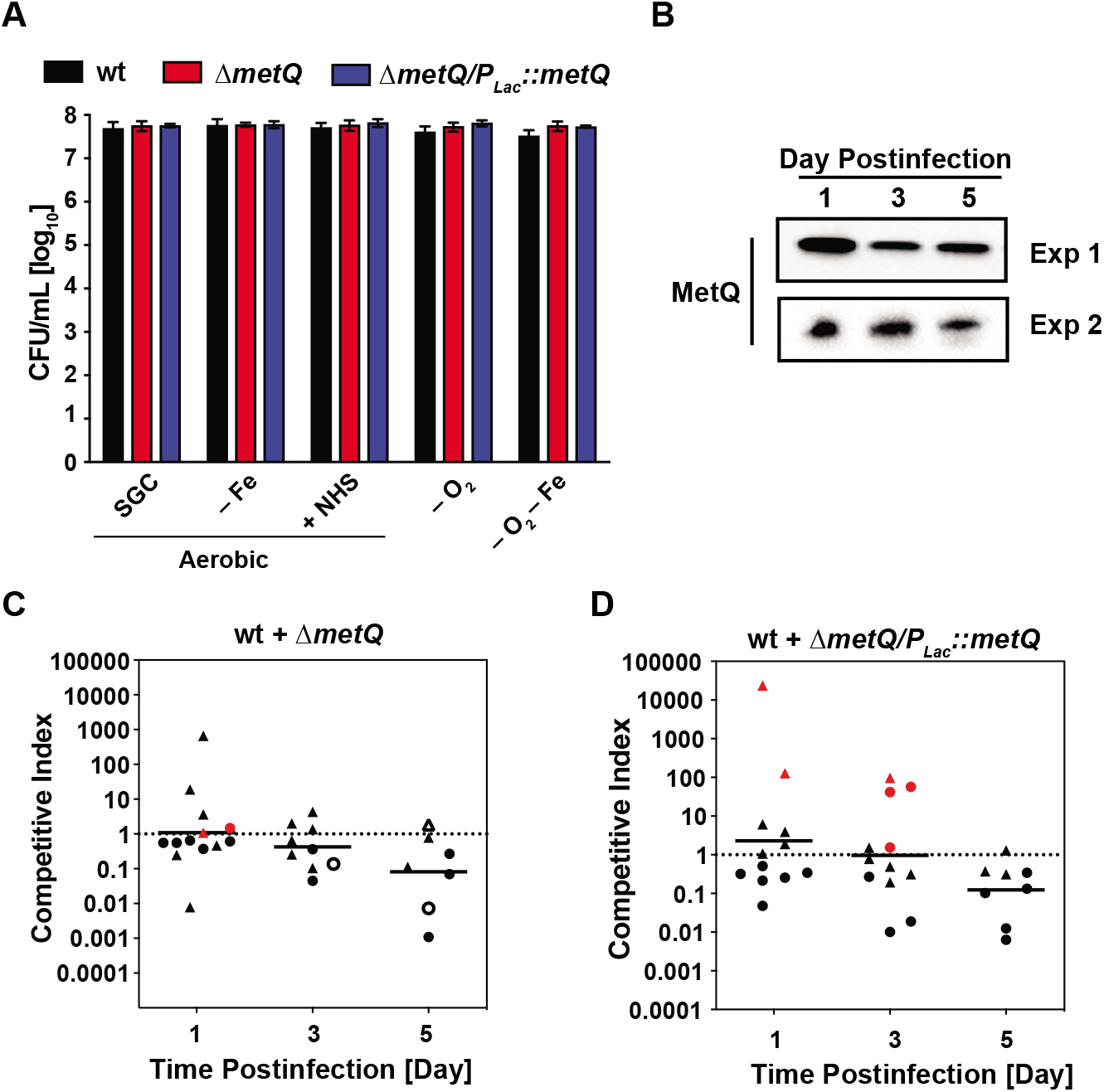
MetQ is expressed during *Ng* infection in the murine lower reproductive tract but is dispensable for bacterial fitness. (**A**) Rapidly growing liquid cultures of strains indicated above the graph were standardized, diluted, and spotted onto solid media for standard growth conditions (SGC), iron limiting conditions (- Fe), presence of normal human serum (NHS), anaerobic conditions (- O_2_), and anaerobic conditions combined with iron limitation (- O_2_ - Fe). CFUs were scored following 22 h incubation. *n* = 3; mean ± SEM. (**B**) Vaginal washes from five female experimentally infected BALB/c mice inoculated with WT *Ng* FA1090 were collected on days 1, 3, and 5 post-inoculation in two independent experiments and pooled washes from each experiment were probed with anti-MetQ antiserum. The amount of sample loaded onto SDS-PAGE was normalized based on the number of gonococcal CFUs recovered from the washes. The relative intensities of MetQ abundance were compared to the amount of MetQ on day 1 post-infection, which was arbitrarily set to 1. (**C**, **D**) Competitive infections between WT bacteria and either the Δ*metQ* mutant (**C**) or the complementation strain Δ*metQ/P_lac_::metQ* (**D**) were performed by inoculating female BALB/c mice intravaginally with approximately equal numbers of each strain (~10^6^ CFU/dose). Vaginal swabs were collected on days 1, 3, and 5 post-infection and enumerated on medium containing streptomycin (total bacteria) or streptomycin and kanamycin (mutant bacteria). The competitive index (CI) was calculated as described in the main text. Experiments were performed on two separate occasions with six mice per group, and the geometric mean of the CI is presented. The assay’s limit of detection of 1 CFU was assigned for any strain not recovered from an infected mouse. Statistical analysis was executed using Kruskal-Wallis and Dunn’s multiple comparison tests to compare statistical significance of CIs between Δ*metQ*/WT and Δ*metQ/P_lac_::metQ*/WT competitions. Red symbols designate mice from which no WT bacteria were recovered, while open symbols signify that no mutant CFU were recovered.

Our prior studies demonstrated that MetQ is ubiquitously expressed in *vitro* [13, 37], however, its cellular pools during infection have not been investigated. Therefore, we next infected female BALB/c mice with WT FA1090 and collected vaginal washes at days 1, 3, and 5 post-infection in biological duplicate experiments. Samples containing equal numbers of viable *Ng* bacteria were separated by SDS-PAGE and probed with anti-MetQ antisera. MetQ was readily detectable at each point examined during the infection period (Fig. 2b). Densitometry analyses using MetQ abundance on day 1 as a reference showed that MetQ levels varied slightly on day 3 post-infection (0.91 ± 0.41; mean ± SEM) and lowered to 0.76-fold (± 0.14; SEM) on day 5. Our investigation indicates that MetQ expression *in vivo* is relatively stable during experimental infection.

Finally, to assess whether MetQ provides *Ng* with a fitness advantage during experimental murine infection, we performed competitive infection experiments in which mice were inoculated vaginally with similar numbers of WT FA1090 mixed with either the Δ*metQ* mutant or the Δ*metQ/P_lac_::metQ* complementation strain. The calculated competitive indices (CIs) for Δ*metQ*/WT were 1.09, 0.42, and 0.08 (geometric means of biological duplicate experiments) on day 1, 3, and 5 post-infection, respectively (Fig. 2c). A similar decrease in CI over time was observed for the Δ*metQ/P_lac_::metQ/WT* competition (Fig. 2 d). Statistical analysis of the data revealed no significant differences. Calculated *p* values for competition experiments conducted *in vitro* were also not significant (Fig. S3).

Based on these studies, we conclude that MetQ is produced *in vivo* but does not provide a growth advantage *in vitro* or a growth or survival advantage during mucosal infection in the lower genital tract of female mice.

### Rationale for immunization and challenge study design

To appraise MetQ as a component of gonorrhea subunit vaccine, we designed a recombinant protein construct using the most prevalent allele (Table S1), by replacing the MetQ lipoprotein signal peptide with an N-terminal 6×histidine tag (Fig. 3a). A highly pure untagged rMetQ antigen that migrated on SDS-PAGE with a predicted molecular weight of 29.87 kDa, which corresponds to mature MetQ, was obtained after two-step chromatography followed by Tobacco Etch Virus (TEV) protease cleavage (Fig. 3b).

**Figure 3.**
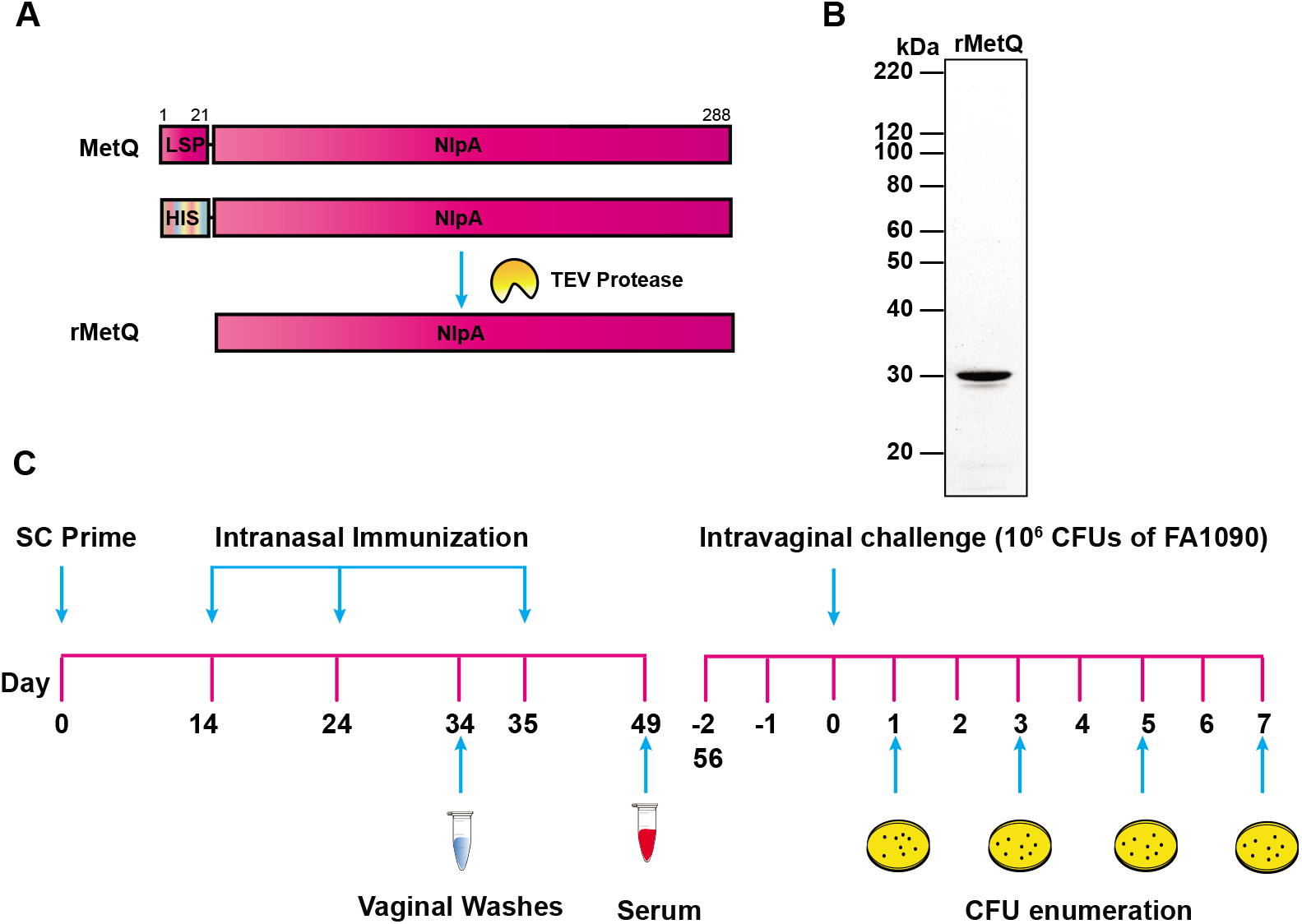
Experimental design of immunization/challenge experiments and MetQ antigen design. (**A**) To generate rMetQ antigen, the *metQ* gene, encoding a full-length MetQ protein, was engineered to produce a recombinant protein that lacks the signal sequence and carries an N-terminal 6×His-tag followed by a Tobacco Etch Virus (TEV) protease cleavage site. (**B**) Soluble rMetQ was purified to homogeneity through several chromatography steps and migrated on SDS-PAGE at approximately 29.87 kDa, consistent with the predicted molecular mass of mature MetQ, as revealed by SYPRO Ruby staining. Untagged rMetQ was used in immunization studies as shown in panel C. (**C**) Female BALB/c mice were randomized into three experimental groups (*n* = 20/group) and given rMetQ-CpG, CpG, or PBS (unimmunized) subcutaneously on day 0, followed by three nasal boosts on days 14, 24 and 35. Vaginal washes were collected 10 days after the second immunization (d34) and serum was collected after the final immunization (day 49) to avoid disruption of the vaginal microenvironment prior to bacterial challenge. Three weeks after the final immunization (d 56 or d-2), only mice that entered into the diestrus or anestrus stage (*n* - number of animals indicated) were treated with 17β-estradiol and antibiotics and challenged with 10^6^ CFU of *Ng* FA1090 two days later (day 0). Vaginal swabs were quantitatively cultured for *Ng* on days 1, 3, 5, and 7 post-bacterial inoculation.

We formulated rMetQ with CpG as an adjuvant based on the limited number of *in vivo* protection studies conducted, which suggest a Th1response is associated with accelerated *Ng* clearance [44–48]. Initially, we conducted a pilot study by immunizing groups of 5 BALB/c mice with rMetQ, rMetQ-CpG, CpG, or PBS using a subcutaneous prime injection, followed by a regimen of three intranasal boost doses. A robust, single protein band corresponding to the molecular weight of rMetQ (~30 kDa) was detected in immunoblots with serum from mice immunized with rMetQ alone (Fig. S4). Immune recognition was only slightly enhanced when antisera to rMetQ-CpG were used (44,405 versus 49,893 relative intensity units, respectively), indicating that rMetQ is a strong immunogen on its own. No signal was detected in sera obtained from animals immunized with adjuvant alone or PBS (Fig. S4). Based on these encouraging results, we performed two large-scale immunization/challenge experiments. To ensure sufficient statistical power for preclinical vaccine research, cohorts (*n*=20/group) of BALB/c mice were given rMetQ formulated with CpG, adjuvant alone, or PBS and challenged vaginally with wild type FA1090 *Ng* three weeks after the final immunization as described [38, 39]. Bacterial clearance was assessed by enumerating *Ng* CFUs present in vaginal washes on days 1, 3, 5, and 7 post-infection (Fig. 3c). Infection duration and bacterial burden were compared between immunized and control groups (Fig. S5). To evaluate the antigen-specific immune responses elicited, vaginal washes and sera collected after the second boost and 14 days after the third boost, respectively, were subjected to immunoblotting and enzyme-linked immunosorbent assays, ELISA (Figs. 5–6).

### Immunization with rMetQ-CpG accelerates clearance from experimentally challenged mice

In the first immunization/challenge experiment, faster clearance was observed in the rMetQ-CpG cohort compared to the CpG- and unimmunized-groups (*p*=0.004 and *p*<0.0001, respectively; Fig. S5a). Mice that received rMetQ-CpG had a 4- and 10-fold lower AUC than mice given either CpG or PBS (*p*>0.9 and *p*=0.0006, respectively; Fig. S5c). In the repeat experiment, the percentage of culture-positive mice over time showed that rMetQ-CpG-immunized mice had a significantly faster clearance rate compared to mice given rMetQ (*p*=0.02) or PBS (*p*=0.001) but not adjuvant alone (Fig. S5d). An area under the curve (AUC) analysis showed that the cumulative burden of infection over time was significantly lower in rMetQ-CpG-immunized mice compared to the rMetQ-, CpG- and PBS-groups (*p*=0.005, *p*=0.03 and *p*=0.003, respectively; Fig. S5f). A cohort of mice given rMetQ showed no difference in the clearance rate or bioburden compared to control groups (Fig. S5 d-f), from which we conclude that formulation with CpG is required for MetQ to confer protection.

The combined data for the two independent MetQ immunization/challenge experiments are shown in Fig. 4. Mice immunized with rMetQ-CpG cleared the infections significantly faster than those given PBS (*p*<0.0001) or adjuvant alone (*p*=0.002; Fig.4a). Importantly, by day 5 and 7 post-challenge, 72.8 and 90.9% of mice that received rMetQ-CpG cleared gonorrhea infection, compared to 42.8 and 65.7% for CpG-inoculated animals and 20.6 and 35.3% in the placebo group, respectively. The gonococcal cumulative burden was also considerably lower in rMetQ-CpG immunized mice in comparison to mice given either CpG or PBS (*p*=0.03 and *p*<0.0001, respectively; Fig. 4b, c).

**Figure 4.**
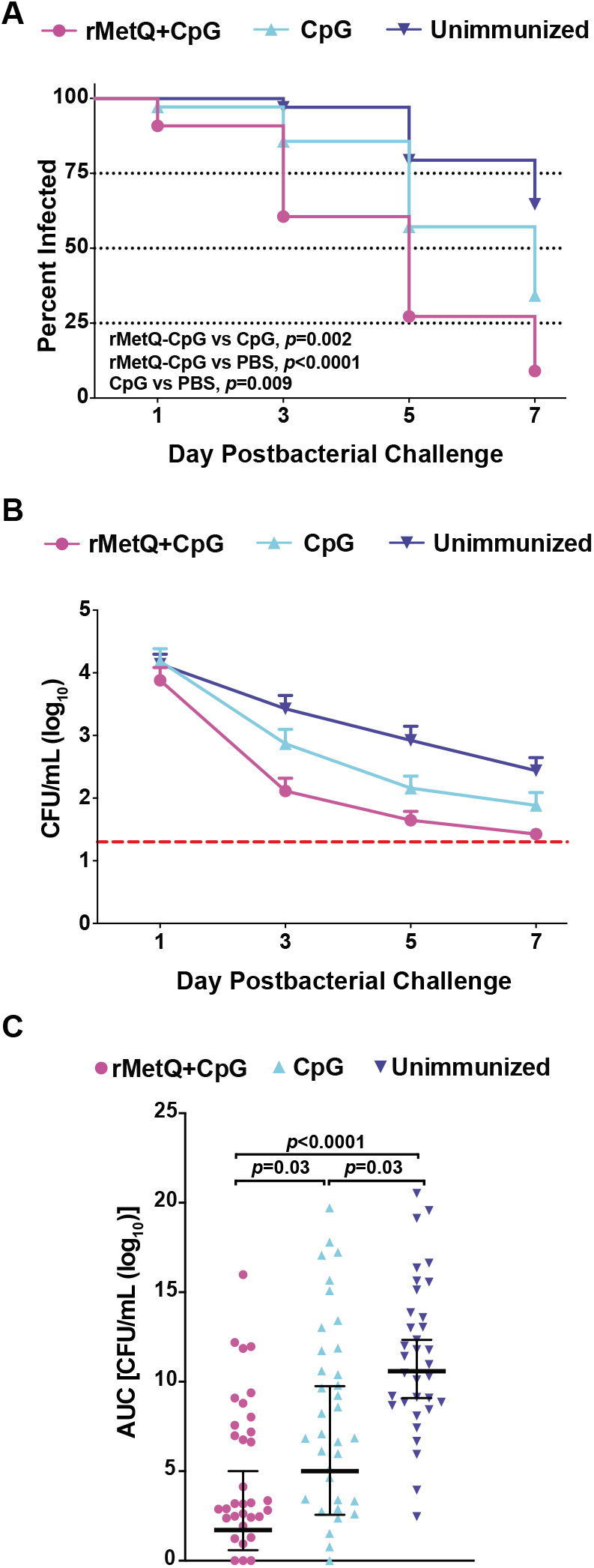
Mice immunized with rMetQ-CpG clear infections significantly faster following vaginal challenge with *Ng*. Groups of BALB/c mice were immunized with rMetQ-CpG or given CpG (adjuvant) alone or PBS (unimmunized) as per the immunization regimen shown in Fig. 3. Subsequently, mice in the diestrus stage or in anestrus were treated with 17β-estradiol and antibiotics and challenged with 10^6^ CFU of strain FA1090 three weeks after the final immunization (*n*=16-18 mice/group). Vaginal washes were quantitatively cultured for *N. gonorrhoeae* on days 1, 3, 5 and 7 post-bacterial challenge. **(A)** The percentage of mice with positive vaginal cultures was plotted over time as Kaplan Meier curves and the results analyzed by the Log Rank test. **(B)** The average number of CFU recovered from each experimental group was plotted over time. The limit of detection was 20 CFU/mL of vaginal swab suspension. This value was used for mice with negative cultures. **(C)** AUC (log_10_ CFU/mL) analysis of murine colonization. Data are presented for individual mice. Horizontal bars represent the geometric mean of the data with the 95% confidence interval. Data shown are combined data from two independent experiments.

### rMetQ-CpG elicits a prominent antigen-specific Th1 response

To comparatively assess the specificity and isotype profiles of antibody responses elicited by rMetQ-CpG vaccination, we performed immunoblotting and ELISA using serum and vaginal washes derived from all cohorts of animals (Figs. 5–6). The immunization regimen induced a strong MetQ-specific serum and vaginal IgG and IgA response as assessed by immunoblot of *Ng* cell envelopes (CE) and purified rMetQ immunogen (Fig. 5a–d). Serum and vaginal IgG and IgA (Fig. 5a, c and 5b, d; respectively) obtained from mice that received rMetQ-CpG readily recognized a single ~30 kDa protein band in the CE samples that aligned with the rMetQ band, but not serum or vaginal washes from control mice (CpG- or PBS-immunized). Overall, relative MetQ intensities were lower in samples probed with vaginal secretions than with serum (Fig. 5 a-b versus c-d). The densitometry analysis of MetQ protein bands detected with serum IgG/IgA and vaginal IgG/IgA in a crude CE protein fraction in comparison to samples containing purified rMetQ, showed that the relative intensities were 2.94-/2.75- and 1.5-/1.9-fold lower, respectively. Together, these experiments demonstrated that immunization of mice with rMetQ-CpG induces antigen-specific immune responses that are both systemic and, critically, at the genital mucosae and that robust MetQ pools reside in the *Ng* cell envelope.

**Figure 5.**
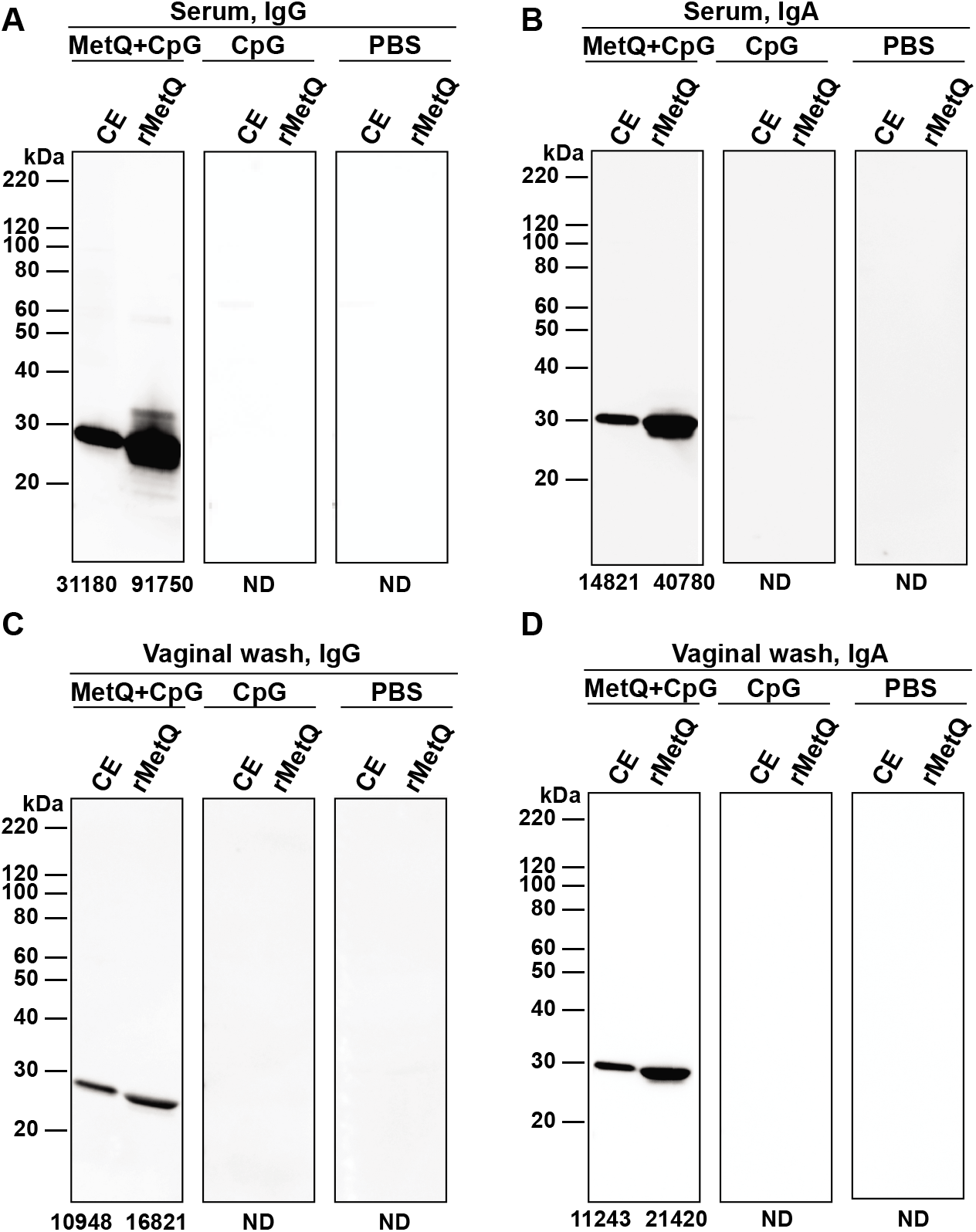
MetQ-specific serum IgG and vaginal IgG and IgA are induced by rMetQ-CpG. Female mice were immunized with rMetQ-CpG, CpG, or PBS in immunization/challenge studies. Total cell envelope (CE) proteins from *Ng* FA1090 and rMetQ were fractionated by SDS-PAGE. Immunoblotting was performed with pooled serum (**A**, **B**) and vaginal washes (**C**, **D**) collected after the third immunization, followed by secondary antibodies against mouse IgG (**A**, **C**) or IgA (**B**, **D**). The intensity of the bands was measured by Fiji software [43] and the value is recorded under each lane. ND-not detected.

**Figure 6.**
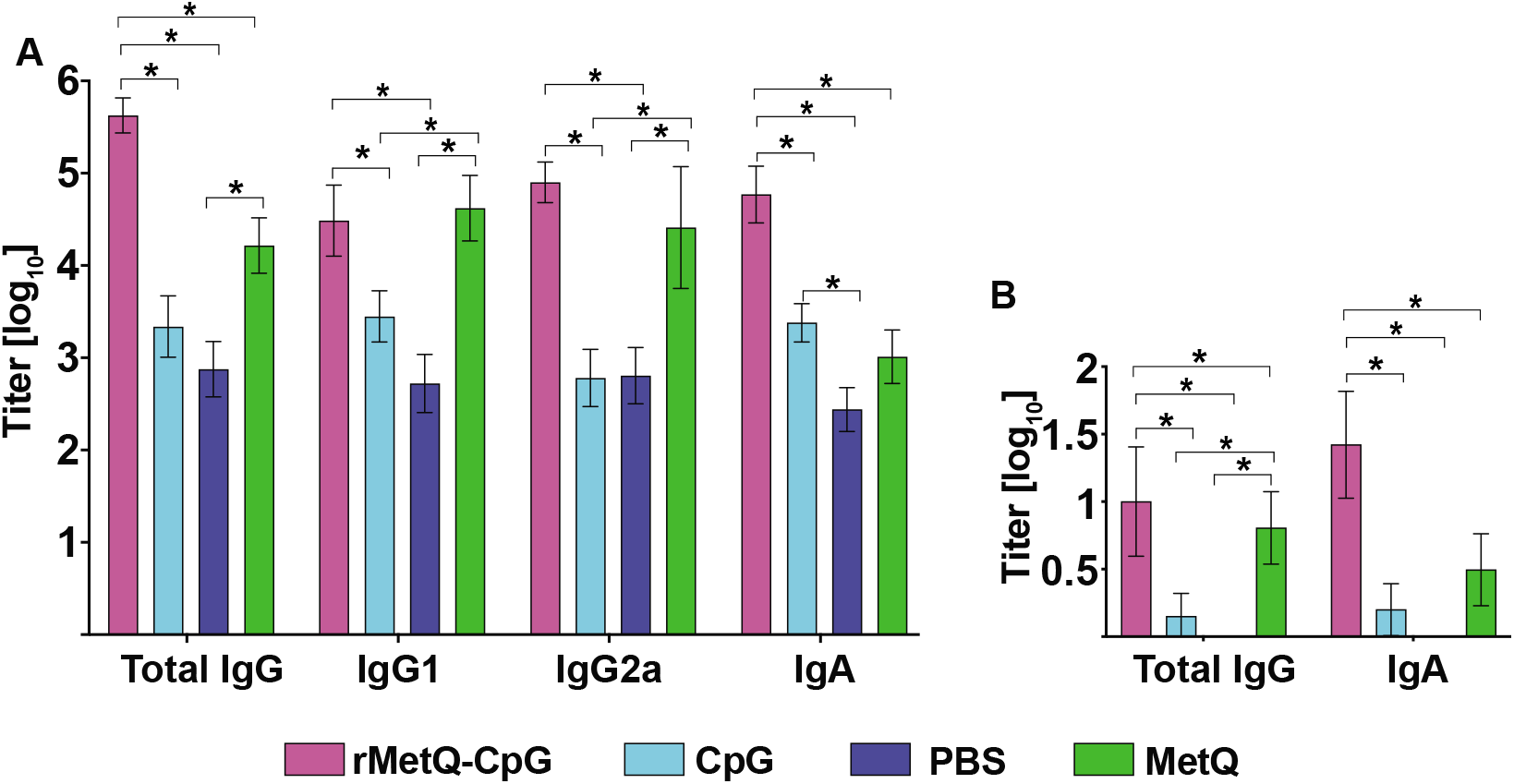
Anti-MetQ antibody titers elicited by rMetQ and rMetQ-CpG immunization. (**A**) Post-immunization (d49) total IgG, IgG1, IgG2a, and IgA antibody titers in sera from mice immunized with rMetQ, rMetQ-CpG, CpG, or unimmunized. (**B**) Post-immunization (d34) vaginal secretions from mice immunized with rMetQ, rMetQ-CpG, CpG, or unimmunized were examined for total IgG and IgA. Bar graphs represent geometric mean ELISA titers with error bars showing 95% confidence limits. Statistical significance between data in groups was determined using Kruskal-Wallis with Dunn’s multiple comparison test; \**p*<0.05.

Analyses of the combined ELISA data showed that total serum IgG and IgA titers in mice immunized with rMetQ-CpG were significantly higher than in mice that received rMetQ alone, adjuvant alone, or PBS (Fig. 6a, Table S3). The calculated geometric mean of total IgG in mice immunized with rMetQ-CpG was 4.2×10^5^ in comparison to 16.5×10^3^, 2.2×10^3^ and 7.5×10^2^ in mice that received rMetQ, CpG, and PBS, respectively. Immunization with rMetQ alone, in comparison to rMetQ-CpG, resulted in similar induction of IgG1 (4.2×10^4^ and 3.1×10^4;^ respectively) and IgG2a antibody responses (2.6×10^4^ and 8.0×10^4;^ respectively) that were significantly greater than in mice immunized with CpG and PBS. However, the IgG1/IgG2a ratio was lower (0.38) in mice given rMetQ formulated with CpG compared to rMetQ alone (1.6), indicating a slight systemic bias toward a Th1 response in the protected group, consistent with the activity of CpG as a Th1-stimulating adjuvant. The serum IgA titers were 57.4-, 24.6-, and 214-fold higher in rMetQ-CpG-immunized mice than in rMetQ, CpG and PBS groups, respectively (Table S3). While minimal IgG and IgA antibody responses were detected in vaginal secretions from unimmunized- and CpG-treated cohorts, rMetQ-CpG vaccination gave significant ~10 and 16-fold increases in IgG (10.02; geometric mean) and IgA (26.42; geometric mean) titers, respectively (Fig. 6b, Table S3). Although immunization with rMetQ alone resulted in a significant increase in MetQ-specific mucosal IgG response (6.3; geometric mean), the levels of secretory IgA were not statistically greater than in control groups. Cumulatively, these analyses demonstrated that rMetQ-CpG formulation elicits significantly high systemic and mucosal antibody titers with a greater Th1 response as predicted by relative serum IgG2a and IgG1 immunoglobulin isotype titers.

We conclude that rMetQ formulated with CpG induces a protective immune response that accelerates *Ng* clearance from the murine lower genital tract, and that while rMetQ alone induces high serum and vaginal antibody responses, rMetQ-CpG increases total IgG and IgA antibodies in serum and vaginal mucosae and skews the immune response towards Th1 response.

## DISCUSSION

In recent years, an increasing number of reports have evaluated the ability of gonorrhea vaccine candidates to elicit immune responses in laboratory animals. However, very few reports examined their capacity to induce protective responses that would enhance clearance and shorten the duration of infection [18]. Accordingly, with this study, we have examined the immune response and protective capability of a lipoprotein antigen discovered through reverse vaccinology: MetQ [NGO2139; [13, 14, 28, 30]]. In light of the extensive surface protein variability inherent to *Ng*, highly conserved antigens should be selected to provide broad vaccine coverage. Our in-depth bioinformatics illustrates that MetQ is a desirable antigen based on its conservation: the MetQ amino acid sequence is identical in nearly 97% of *Ng* isolates globally (Fig. 1, Table S1). Among the 17 amino acid polymorphisms, single changes are found in 0.02-2.7 % of isolates and a single allele with 8 divergent amino acids is present in only 0.05% of isolates. Consequently, vaccination with the single most prevalent MetQ variant will likely provide broad protection against the majority of *Ng* encountered worldwide. Additionally, prior analysis of the *Nm* homolog, GNA1946, in 97 isolates revealed the existence of only 20 different alleles with 23 and 5 polymorphic nucleotide and amino acid sites, respectively [49]. In combination with our bioinformatic analyses (Figs. S1 and S2), conservation data from *Nm* suggest that MetQ may be a suitable antigen for a universal *Neisseria* vaccine. Most importantly our studies showed that rMetQ-CpG vaccination both significantly accelerated clearance and reduced bacterial burden in infected animals (Fig. 4). Although, anti-MetQ antibodies block protein function and are bactericidal [13, 14],

We found that the addition of CpG was critical to promote a protective response with the rMetQ immunogen (Fig. 4 and Fig. S5). CpG oligodeoxynucleotides resemble bacterial DNA and provoke an immune response. Their mechanism of action relies on the stimulation of Toll-like receptor (TLR) 9, which is primarily expressed in human B cells and plasmacytoid dendritic cells (pDCs). Upon CpG-induced TLR 9 activation, both B cells and pDCs upregulate expression of major histocompatibility complex (MHC) class II molecules and release cytokines that signal naïve T cells to differentiate to Th1 cells. B cells then differentiate into either plasma cells or memory cells, while pDCs also differentiate to a stage where antigen processing and presentation are enhanced [50]. The ability of CpG to augment antigen presentation through DCs may therefore overcome the antigen presentation restriction *Ng* imposes on macrophages. The multi-target mechanism of CpG action is also consistent with data that indicated Th1 and B cell responses were both important for immune memory against gonorrhea. Wild type mice administered IL-12 during infection were able to clear repeat infections more quickly than untreated mice. In contrast, B cell-deficient mice were impaired in their ability to clear the repeat infection [47]. In our experiments, mice cohorts that received rMetQ-CpG developed a robust, MetQ-specific antibody responses in both serum and vaginal secretions (Figs. 5, 6). The presence of antigen-specific IgA and IgG in the vaginal mucosae is promising, as it suggests that vaccination would result in an antibody response at the site of infection. Together, these data suggest the induction of IgA and a Th1 responses are likely to be important for effective gonococcal clearance. This conclusion is supported by studies with experimental vaccines formulated with Th1-inducing adjuvants or delivery systems [44–48].

The data presented here warrant further development of rMetQ as a gonorrhea vaccine candidate. Subunit vaccines are attractive for their safety, stability, and defined composition. One of the two available licensed serogroup B *Nm* vaccines, Trumenba (Pfizer), is a subunit vaccine that contains two factor H binding protein variants, which were identified through reverse vaccinology [51]. Although Trumenba demonstrates that a single-component subunit vaccine can be successful, to provide superior immune responses against the sophisticated *Ng*, a cocktail vaccine with several other highly conserved surface antigens is likely needed. The current renaissance in the gonorrhea vaccine field brings optimism for generating a much needed, effective gonorrhea vaccine by advancing vaccine workflows to preclinical and clinical trials, discovering conserved immunogens, and illuminating new concepts to counteract the ways *Ng* subverts the immune system.

## ACKNOWLEDGEMENTS

We thank Dr. Kristi Conolly (Uniformed Services University of the Health Sciences) for helpful discussion regarding immunization/challenge experiments.

## SUPPORTING INFORMATION

**Supplemental Figure S1. *metQ* nucleotide variation across *Neisseria* genus**.

**Supplemental Figure S2. Phylogenetic analysis of MetQ variation among *Neisseria***.

**Supplemental Figure S3. *In vitro* competition assay**.

**Supplemental Figure S4. Immunization with rMetQ alone or with CpG induces antigen-specific immune response**.

**Supplemental Figure S5**. Infection dynamics from individual rMetQ-CpG immunization experiments. In the first immunization/challenge experiments cohorts of BALB/c mice were immunized with rMetQ-CpG, CpG, or PBS (A, C, E). The second experiments consisted of groups that were given rMetQ, rMetQ-CpG, CpG, or PBS (B, D, F) as per the immunization regimen shown in Fig. 3. Mice in the diestrus stage or in anestrus were treated with 17β-estradiol and antibiotics and challenged with 10^6^ CFU of strain FA1090 three weeks after the final immunization. Vaginal washes were quantitatively cultured for *Ng* on days 1, 3, 5 and 7 post-bacterial challenge. **(A, B)** The percentage of mice with positive vaginal cultures was plotted over time as Kaplan Meier curves and the results analyzed by the Log Rank test. **(C, D)** The average number of CFU recovered from each experimental group was plotted over time. The limit of detection was 20 CFU/mL of vaginal swab suspension. This value was used for mice with negative cultures. **(E, F)** AUC (log 10 CFU/mL) analysis of murine colonization. Data are presented for individual mice. Horizontal bars represent the geometric mean of the data with the 95% confidence interval. Data shown are combined data from two independent experiments.

**Supplemental Table S1. Analysis of MetQ alleles and number of *Ng* isolates per allele grouping**.

**Supplemental Table S2. Etest antibiotic susceptibility experiments**.

**Supplemental Table S3. Geometric means antibody titers measured in serum and vaginal washes on days 34 and 49, respectively, since the SC immunization**.

Geometric mean titers of antibodies against purified rMetQ were determined as described in Materials and Methods.

**Supplemental Movie S1**. *Ng* amino acid polymorphisms mapped to *Nm* MetQ crystal structure bound to L-methionine, rotated about the X axis. Movie generated in PyMol.

**Supplemental Movie S2**. *Ng* amino acid polymorphisms mapped to *Nm* MetQ crystal structure bound to L-methionine, rotated about the Y axis. Movie generated in PyMol.

**Supplemental Movie S3**. *Ng* amino acid polymorphisms mapped to *Nm* MetQ crystal structure in the substrate-free conformation, rotated about the X axis. Movie generated in PyMol.

**Supplemental Movie S4**. *Ng* amino acid polymorphisms mapped to *Nm* MetQ crystal structure in the substrate-free conformation, rotated about the Y axis. Movie generated in PyMol.

