## Supplemental Information for "A novel gonorrhea vaccine composed of MetQ lipoprotein formulated with CpG shortens experimental murine infection"

##### Fig S1

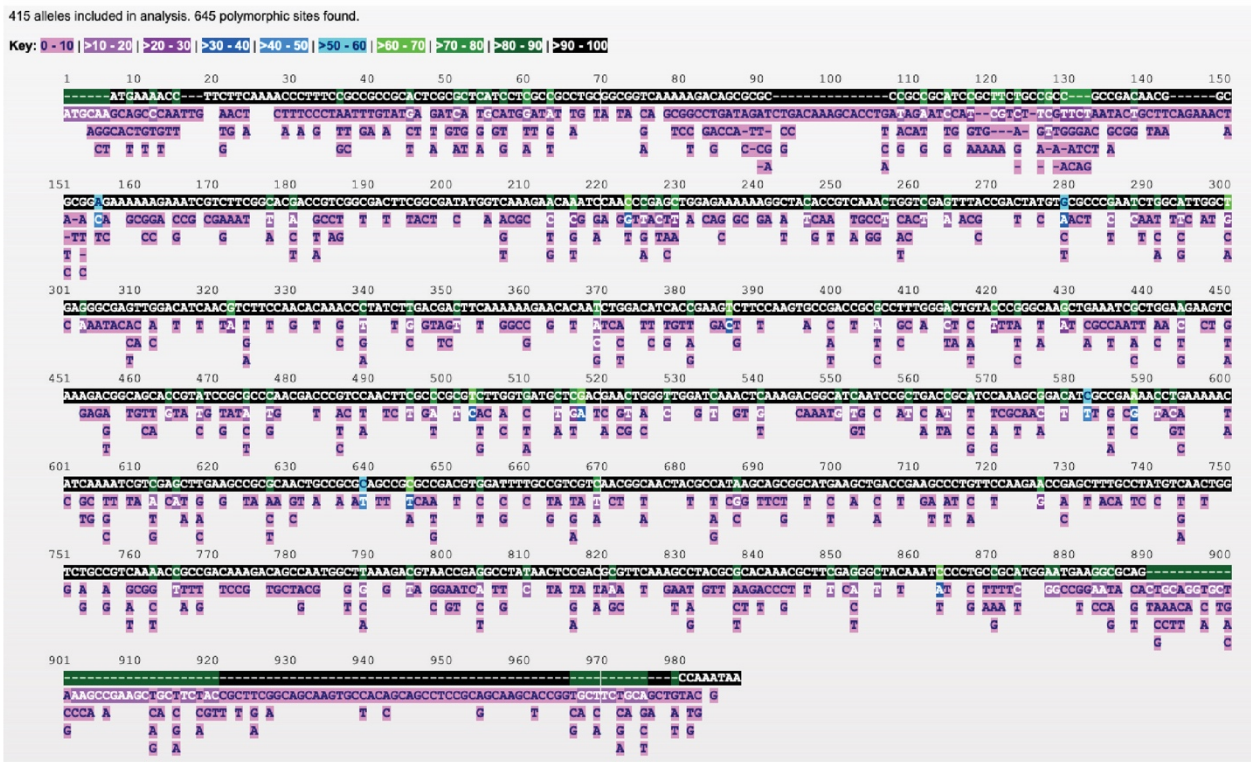


**Supplemental Figure S1. *metQ* nucleotide variation across *Neisseria* genus.** Nucleotide polymorphic sites in the *metQ* locus (NEIS1917) of 21,798 *Neisseria* isolates with *metQ* sequence data deposited into the Neisseria PubMLST database.

**Fig S2**

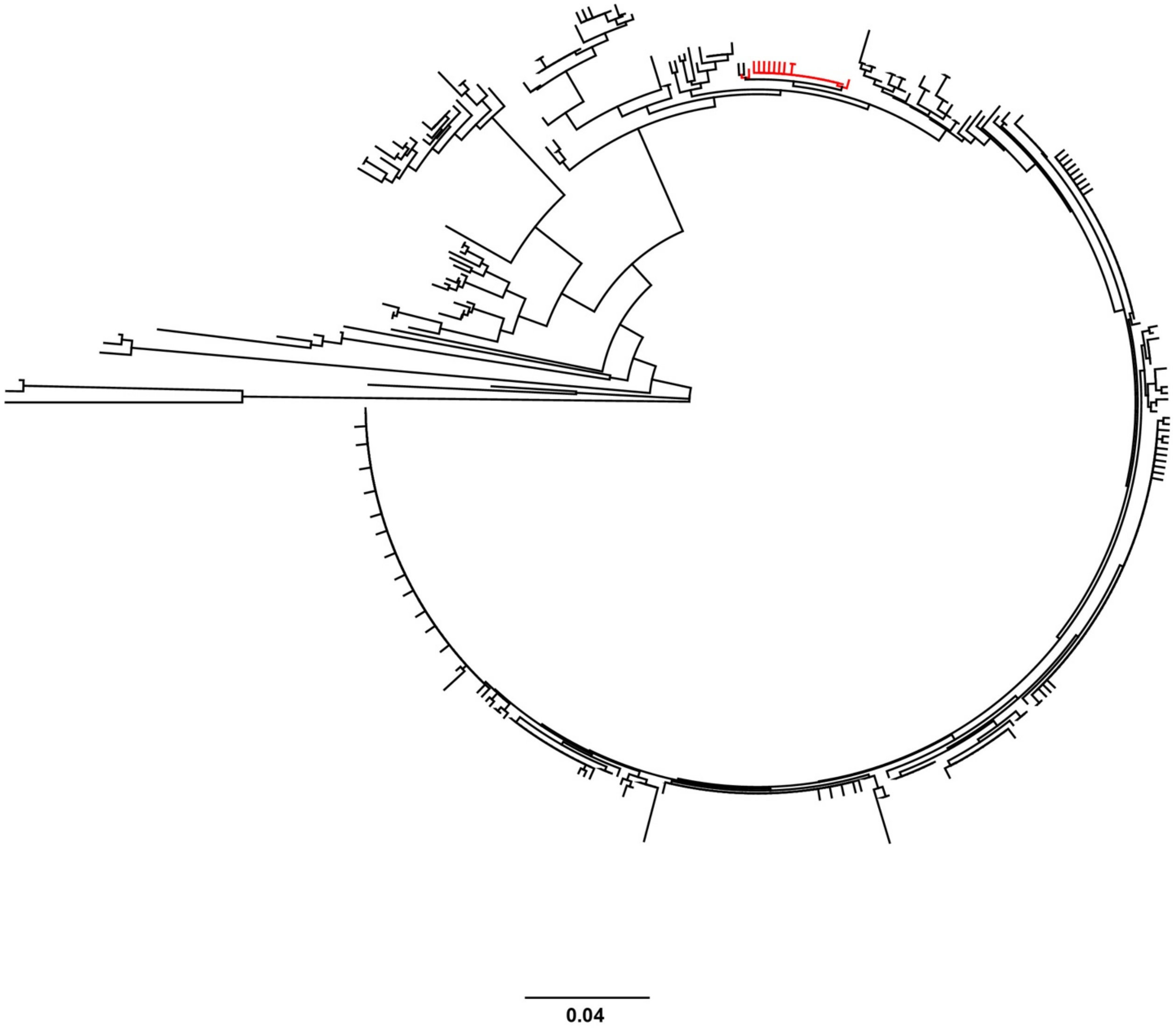


**Supplemental Figure S2. Phylogenetic analysis of MetQ variation among *Neisseria*.** A Neighbor-Joining tree was constructed in Geneious using the Jukes-Cantor genetic distance model for translations of all *metQ* nucleotide sequences in the *Neisseria* PubMLST database. Branches of *Ng* alleles are highlighted in red.

### Fig S3

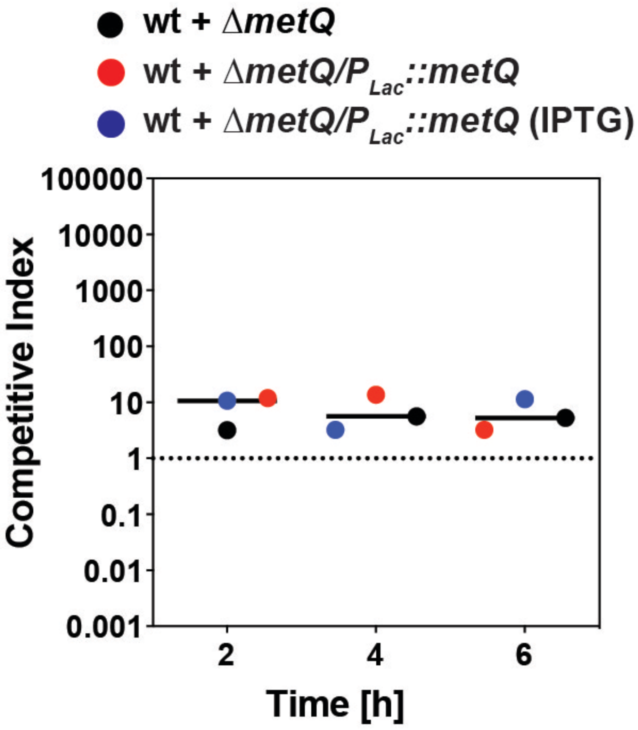


**Supplemental Figure S3. *In vitro* competition assay.** WT bacteria were combined with equal numbers of either Δ*metQ* bacteria or the Δ*metQ*/$P_{lac}$::*metQ* complementation strain and propagated in liquid medium. Competitions with the complementation strain were performed in the presence and absence of IPTG. At timepoints indicated, cultures were enumerated and the competitive index was calculated.

#### Fig. S4

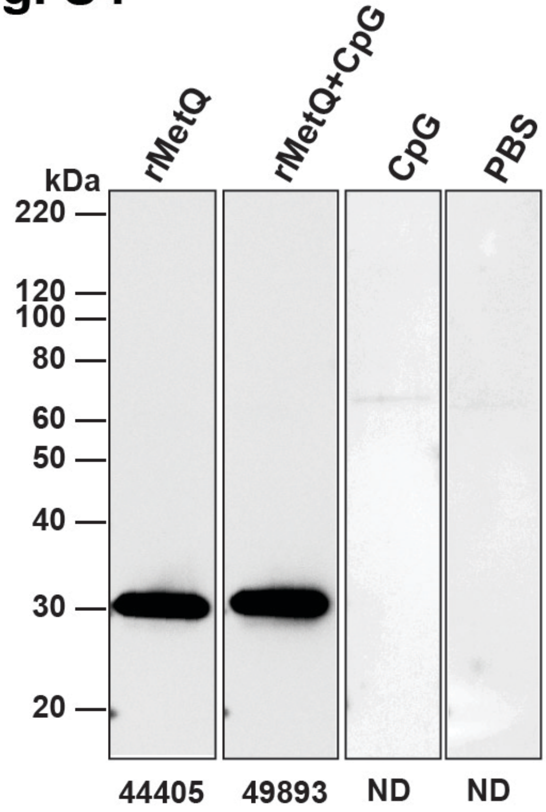


**Supplemental Figure S4. Immunization with rMetQ alone or with CpG induces a specific immune response.** Female mice were immunized with rMetQ, rMetQ-CpG, CpG, or PBS in the pilot immunization study. Total cell envelope proteins from *Ng* FA1090 were separated by SDS-PAGE. Immunoblotting was performed with pooled serum collected after the third immunization. The band intensity was measured by Fiji software and the value is recorded under each lane. ND-not detected.

Fig S5

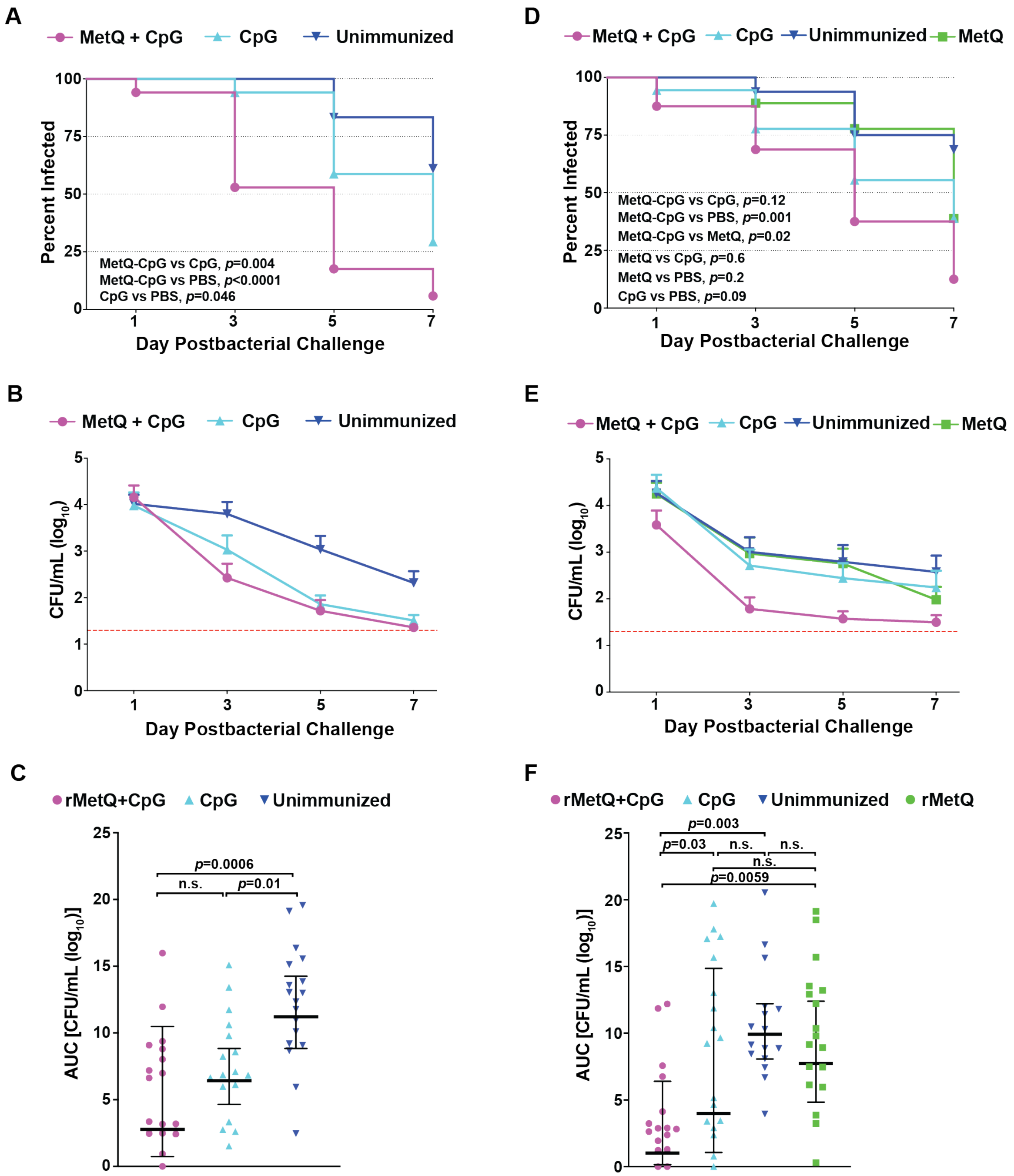


**Supplemental Figure S5.** Infection dynamics from individual rMetQ-CpG immunization experiments. In the first immunization/challenge experiments cohorts of BALB/c mice were immunized with rMetQ-CpG, CpG, or PBS (A, C, E). The second experiments consisted of groups that were given rMetQ, rMetQ-CpG, CpG, or PBS (B, D, F) as per the immunization regimen shown in Fig. 3. Mice in the diestrus stage or in anestrus were treated with 17β-estradiol and antibiotics and challenged with $10^6$ CFU of strain FA1090 three weeks after the final immunization. Vaginal washes were quantitatively cultured for *Ng* on days 1, 3, 5 and 7 post-bacterial challenge. **(A, B)** The percentage of mice with positive vaginal cultures was plotted over time as Kaplan Meier curves and the results analyzed by the Log Rank test. **(C, D)** The average number of CFU recovered from each experimental group was plotted over time. The limit of detection was 20 CFU/mL of vaginal swab suspension. This value was used for mice with negative cultures. **(E, F)** AUC ($\log_{10}$ CFU/mL) analysis of murine colonization. Data are presented for individual mice. Horizontal bars represent the geometric mean of the data with the 95% confidence interval. Data shown are combined data from two independent experiments.

**Supplemental Table S1. Analysis of MetQ alleles and number of *Ng* isolates per allele grouping.** All *Ng* nucleic acid *metQ* alleles present in the *Neisseria* PubMLST database as of March 26, 2020, sorted by prevalence. Percentages calculated according to the total number of *Ng* isolates with *metQ* sequence information deposited into the database. Translations of the nucleic acid sequences were aligned and the amino acid sequences were compared to the most common allele (10) to assess overall amino acid conservation.

| Allele | # of *Ng* Isolates | % of Total *Ng* Isolates (*n* = 4,421) | # of Divergent AAs compared to allele 10 |
|---|---|---|---|
| 10 | 2238 | 50.62% | 0 |
| 8 | 1055 | 23.86% | 0 |
| 171 | 569 | 12.87% | 0 |
| 45 | 379 | 8.57% | 0 |
| 266 | 115 | 2.60% | 1 |
| 246 | 25 | 0.57% | 0 |
| 608 | 8 | 0.18% | 0 |
| 250 | 5 | 0.11% | 1 |
| 248 | 4 | 0.09% | 0 |
| 252 | 4 | 0.09% | 1 |
| 172 | 3 | 0.07% | 1 |
| 41 | 2 | 0.05% | 8 (including 1 gap) |
| 191 | 2 | 0.05% | 0 |
| 249 | 2 | 0.05% | 1 |
| 253 | 2 | 0.05% | 1 |
| 475 | 2 | 0.05% | 1 |
| 247 | 1 | 0.02% | 1 |
| 251 | 1 | 0.02% | 1 |
| 294 | 1 | 0.02% | 0 |
| 437 | 1 | 0.02% | 0 |
| 473 | 1 | 0.02% | 1 |
| 474 | 1 | 0.02% | 1 |

**Supplemental Table S2. Etest antibiotic susceptibility experiments.**

Minimal inhibitory concentrations, evaluated by Etest antimicrobial test strips. [a]MIC values are presented in µg/mL. [b]Vector for MetQ complementation encodes erythromycin resistance, which also provides resistance against azithromycin.

| | **WT**[a] | **∆*metQ***[a] | **∆*metQ*/P$_{lac}$::*metQ***[a] |
|---|---|---|---|
| **Polymyxin B** | 64 | 64 | 64 |
| **Vancomycin** | 8 | 8 | 8 |
| **Azithromycin** | 0.032 | 0.032 | 0.5[b] |
| **Cefotaxime** | 0.004 | 0.004 | 0.008 |
| **Ampicillin** | 0.125 | 0.125 | 0.125 |
| **Tetracycline** | 0.125 | 0.125 | 0.125 |
| **Benzylpenicillin** | 0.064 | 0.064 | 0.064 |

**Supplemental Table S3. Geometric means antibody titers measured in serum and vaginal washes on days 34 and 49, respectively, since the SC immunization.**

Geometric mean titers of antibodies against purified rMetQ were determined as described in Materials and Methods.

| | rMetQ-CpG | CpG | PBS | rMetQ |
|---|---|---|---|---|
| **Serum** | | | | |
| **Total IgG** | 422876 | 2187 | 752.2 | 16548 |
| **IgG1** | 30727 | 2811 | 527.7 | 41890 |
| **IgG2a** | 80121 | 603.9 | 643.0 | 25904 |
| **IgA** | 59049 | 2403 | 275.5 | 1028 |
| **Vaginal wash** | | | | |
| **Total IgG** | 10.02 | 1.41 | 1.00 | 6.38 |
| **IgA** | 26.42 | 1.58 | 1.00 | 3.12 |

**Supplemental Movie S1.** *Ng* amino acid polymorphisms mapped to *Nm* MetQ crystal structure purified under standard conditions, rotated about the X axis. Movie generated in PyMol.

**Supplemental Movie S2.** *Ng* amino acid polymorphisms mapped to *Nm* MetQ crystal structure purified under standard conditions, rotated about the Y axis. Movie generated in PyMol.

**Supplemental Movie S3.** *Ng* amino acid polymorphisms mapped to *Nm* MetQ crystal structure purified under D-methionine conditions, rotated about the X axis. Movie generated in PyMol.

**Supplemental Movie S4.** *Ng* amino acid polymorphisms mapped to *Nm* MetQ crystal structure purified under D-methionine conditions, rotated about the Y axis. Movie generated in PyMol.
